# SCS313 Serrano, a big trefoil selected for flowering ability and seed production

**DOI:** 10.1101/2021.10.26.465940

**Authors:** Dediel A. Rocha, Ulisses A. Cordova, Jefferson Flaresso, Joseli Stradiotto, Murilo Dalla-Costa

## Abstract

SCS313 Serrano is a tetraploid cultivar of *Lotus uliginosus* developed by the Research and Rural Extension Company of Santa Catarina, Brazil, to improve flowering ability and seed production in low-latitude regions. SCS313 Serrano was developed from the selection of individual plants of the cultivar Grasslands Maku. Seeds from the initial breeding population was bulked and used to establish a field trial with spaced plants. The plants were selected and classified as late or early, regarding the beginning of flowering, through visual evaluations based on the time for the beginning of flowering. Selected plants were transplanted to crossing blocks and seed harvested on individual plants; a progeny test was conducted, with four replications, through the establishment of approximately 100 genotypes in a field. This process was repeated for three consecutive cycles of recurrent selection. Considering the mean time for the beginning of flowering and vigor performance, the best five genotypes were selected as parental lines for the synthetic cultivar SCS313 Serrano. A morphological difference between SCS313 Serrano and Grasslands Maku is that SCS313 Serrano has absence of hairs in stems whereas Grasslands Maku has a medium stem hair density. In addition, SCS313 Serrano exhibits profuse flowering ability while Grasslands Maku exhibits very sparse seedhead formation. SCS313 Serrano exhibited good persistence under wet conditions and similar forage yields compared to other lotus commercial cultivars. Thus, SCS313 Serrano is recommended to be used as pasture in mixtures with grass in livestock systems, mainly on wet soils.

## Introduction

SCC313 Serrano (*Lotus uliginosus* Schkuhr) is a tetraploid cultivar derived after hybridizing individual plants from a short day-flowering type, moderate seed yield population, selected in Santa Catarina, Brazil, from the cultivar Grasslands Maku. *L. uliginasus* is commonly known as greater birdsfoot trefoil, or big trefoil; it is it a perennial forage legume native to Europe and North Africa (ARMSTRONG, 1974). Compared to birdsfoot trefoil (*L. corniculatus*), big trefoil has larger leaves, and more vigorous underground stems or rhizomes. Even though the cultivar Grasslands Maku was introduced to Uruguay from New Zealand in the 80’s and had shown good adaptation (INIA, 2021), it had not the same success in Brazil, mainly because of the long photoperiod requirement to produce a profitable quantity of seeds.

In general, Lotus species are long-day species, requiring a minimum day length of 14 hours for flowering and therefore producing seeds in regions at latitudes of about 40–50 (PIANO; PECETTI, 2010). The shorter day length requirements of SCC313 Serrano favors natural reseeding in pastures and enhance sward persistence in low-latitude regions, such as southern Brazil.

Big trefoil is generally used for pastures in mixtures with grasses or other legumes, but it can also be used for hay and silage. Its annual forage production is outstanding, maintaining its production more stable when compared to other lotus species. Big trefoil can produce high-quality forage, is tolerant to grazing, and does not cause bloating. Condensed tannins in big trefoil plants protect proteins from rumen degradation and loss, increasing intestinal flow and absorption at the same level. When big trefoil is used single, it can cause a reduction in voluntary consumption by its impact on ruminal microflora, which reduces fiber digestibility (WAGHORN et al., 1994). Nevertheless, big trefoil produces excellent yields when used in mixtures with others species.

Big trefoil is adapted to the poorly drained soils, tolerates low fertility, and aluminum toxicity in acidic soils (SCOTT; MILLS, 1981). SCC313 Serrano is a potentially useful cultivar for many areas in South America, as well as other regions with extensive livestock enterprises.

## Methods

### Selection and Breeding

A mass selection with 600 plants of the cultivar Grasslands Maku that showed flowering, was carried out in a field of 1.3 ha for the development of initial the breeding population. Seeds of the initial breeding population was bulked and used to establish a field trial with 478 plants, with spacing of 0.6 m within rows and 1 m between rows. Then, 105 plants were selected and classified as late or early, regarding the beginning of flowering, through visual evaluations based on the time for the beginning of flowering.

In 1987, sixteen crossing blocks of 105 plants were transplanted, eight blocks for late and eight blocks for early plants. In the following years, a progeny test was conducted, with four replications, through the establishment of 82 genotypes in the field, under the same spacing of the field trial. The cultivars Grassland Maku and São Gabriel, and other experimental populations were included in the progeny test as controls. Plants in the nursery were individually visually rated for time for the beginning of flowering and vigor. Nurseries were green-chopped to a 5-cm stubble height with approximately 30-day intervals, usually after the visual ratings. Nine genotypes were transplanted to an isolated nursery for producing and harvesting of seeds. This process was repeated for three consecutive cycles of recurrent selection. The best five genotypes out of the 82 evaluated, based on the mean time for the beginning of flowering and vigor performance, over this period, were selected and seed bulked to produce the Syn-1. The Syn-2 generation was then produced in isolation in Lages (Santa Catarina, Brazil), from the Syn-1 seeds; this generation was designated as the prebreeder seed and stored at the Research and Rural Extension Company of Santa Catarina (Epagri). The syn-3 seed was produced from the Syn-2 pre-breeder seeds in Lages in 1994 and was designated as the breeder seed of the cultivar SCC313 Serrano.

### Evaluation

Morphological data were collected from individual plants (60) of the cultivars São Gabriel, Grasslands Maku, and SCC313 Serrano. A randomized complete block experimental design was used with four blocks. Each plot of each entry within each block consisted of a row of 10 plants. Each plant was measured for the following characteristics: central leaflet length, central leaflet width, and longest stem length. Visual assessment of individual plants was carried out to access characteristics based on guidelines, to conduct distinctness, uniformity, and stability tests for big trefoil.

Herbage dry matter yield was assessed in performance trials in Lages, Canoinhas, Campos Novos, and Chapecó, in the state of Santa Catarina, Brazil. A randomized complete blocks experimental design was used, with four replicates. Each experimental unit was composed of eight rows of 4.5 m, spaced 0.20 m apart.

### Characteristics

#### Morphological Characteristics

The goal of the Epagri breeding program was to improve flowering, seedhead, and forage characteristics of the parent cultivar Grasslands Maku via recurrent selection. The cultivars São Gabriel, Grasslands Maku, Trojan, and Larrañaga were used as controls to compare botanical and agronomic characteristics. SCS313 Serrano has larger leaflet width and length than São Gabriel and equal to Grasslands Maku. The height of the longest stem is similar to that of São Gabriel. The 1000-seed weight of São Grabriel is higher than that of SCS313 Serrano (Table1). A morphological difference between SCS131 Serrano and Grasslands Maku is that SCS313 Serrano has no stem hairs, whereas Grasslands Maku has a medium stem hair density (Figure 1). SCS313 Serrano exhibits profuse flowering ability in regions at low latitudes, as shown by its seedhead formation, whereas it is very sparse in the cultivar Grasslands Maku (Figure 2).

**Table 1.** Morphological characteristics of different lotus cultivars. Sixty randomly chosen plants was used per cultivar.

| Cultivar | Central leaflet (mm) |  | Longest stem height (cm) | 1000-seed weight |
| --- | --- | --- | --- | --- |
|  | Length | Width |  |  |
| SCC313 Serrano | 15.4 | 10.8 | 25.7 | 0.7 |
| São Gabriel | 11.3 | 6.8 | 26.9 | 1.2 |
| Grasslands Maku | 19.3 | 14.8 | 26.1 | - |
| LSD* ($\alpha = 0.05$) | 1.3 | 0.9 | 1.7 | |

\*Fisher-LSD bonferroni test

**Figure 1.**
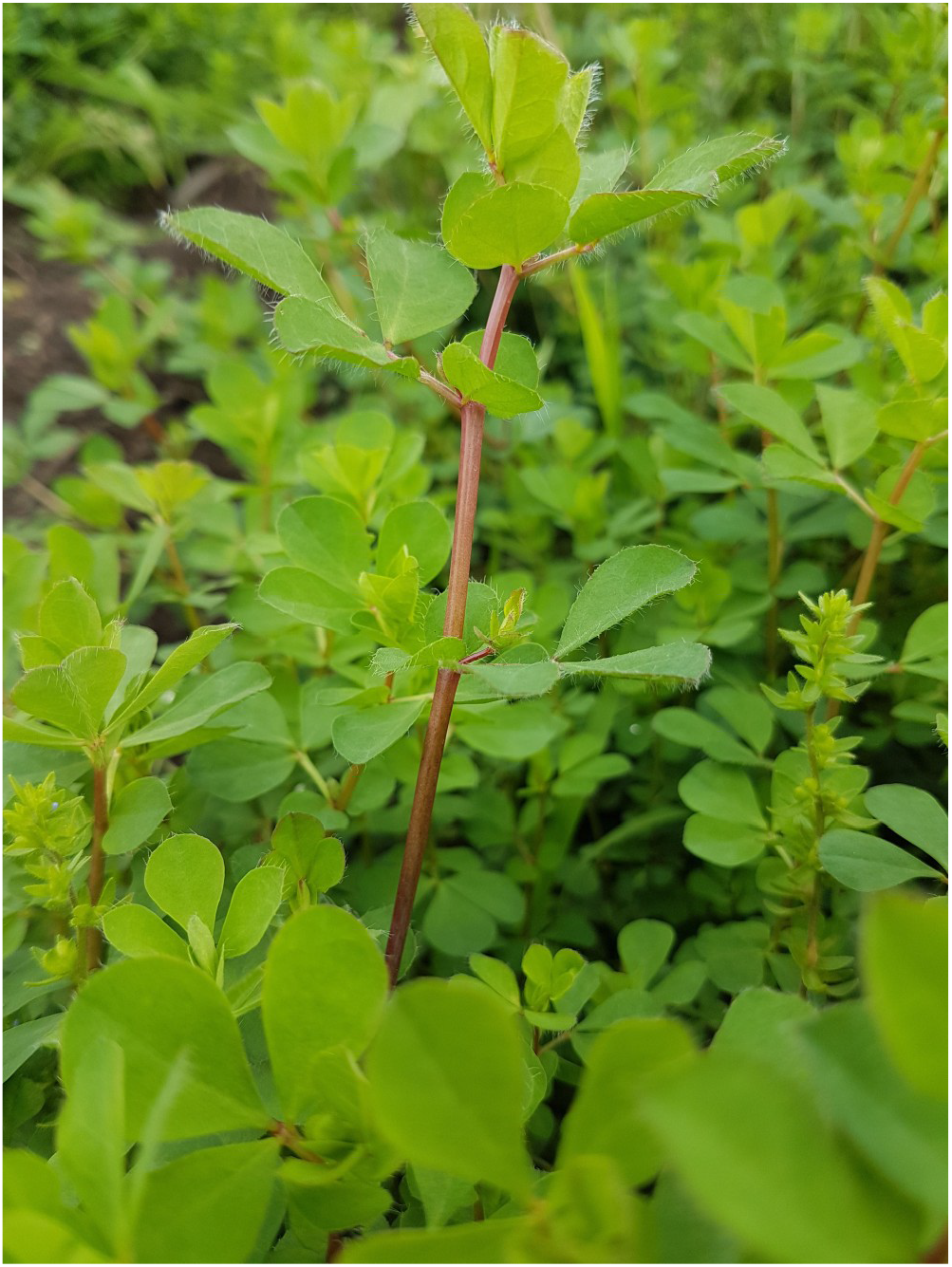
Details of a plant of the cultivar SCS313 Serrano showing leaflets with medium stem hair density; and absent or very sparse stem hair density.

#### Agronomic Performance

The trials conducted across four locations showed that SCS313 Serrano presented similar forage yields to those of the cultivars Trojan and Larrañaga, when it was planted in performance trials (Table 2). The stand survival evaluations in the trials in Lages showed that the stand survival of SCS313 Serrano is lower than that of the cultivar São Gabriel, under prevalent dry condition. It is also shown by the productivity decrease on the third evaluation year (Table 3). However, SCS313 Serrano exhibited good persistence under wet conditions. The seasonally crude protein yield of SCS313 Serrano is equivalent to that of the cultivars Trojan and Larrañaga (Table 4). SCS313 Serrano exhibited less in vitro dry matter digestibility than Larrañaga (Table 5).

**Table 2.** Annual herbage dry matter yields (kg ha^-1^) of different legume cultivars in different testing environments, in Santa Catarina, Brazil, from 2011 to 2012.

| Cultivar | Species | Chapecó |  | Campos Novos |  | Lages |  | Canoinhas |
| --- | --- | --- | --- | --- | --- | --- | --- | --- |
|  |  | 2011 | 2012 | 2011 | 2012 | 2011 | 2012 | 2011 |
| SCS 313 Serrano | *Lotus uliginosus* | 3687 | 5225 | 1906 | 583 | 4565 | 737 | 5013 |
| Trojan | *Lotus uliginosus* | 2975 | 3291 | 2209 | 1377 | 3333 | 981 | 5002 |
| Larrañaga | *Lotus tenuis* | 3100 | 2719 | 7077 | 1809 | 5631 | 3536 | 6731 |
| LSD* ($\alpha = 0.05$) | | ns | 1600 | ns | ns | ns | 1601 | ns |

\*Fisher-LSD bonferroni test

**Table 3.** Annual herbage dry matter yields (kg ha^-1^) of different legume genotypes evaluated for three years in highlands of the state of Santa Catarina, Brazil, from 2018 to 2020.

| Genotype | Species | 2018 | 2019 | 2020 |
| --- | --- | --- | --- | --- |
|  |  | kg.ha-1 |  |  |
| SCS313 Serrano | *Lotus uliginosus* | 3102 c | 9769 a | 2567 b |
| São Gabriel | *Lotus corniculatus* | 6259 a | 9437 a | 6654 a |
| PGGLegP2 | *Lotus corniculatus* | 5893 a | 7840 abc | 5363 ab |
| PGGLegP3 | *Lotus corniculatus* | 5577 ab | 9244 ab | 6095 a |
| PGGLegP1 | *Trifolium repens* | 3026 c | 6296 bc | 4799 ab |
| Zapican | *Trifolium repens* | 3864 bc | 4964 c | 2915 b |

Means followed by the same letters in column are not significantly different from each other by Fisher's Least Significant Difference (LSD) procedure at $P < 0.05$ .

**Table 4.**
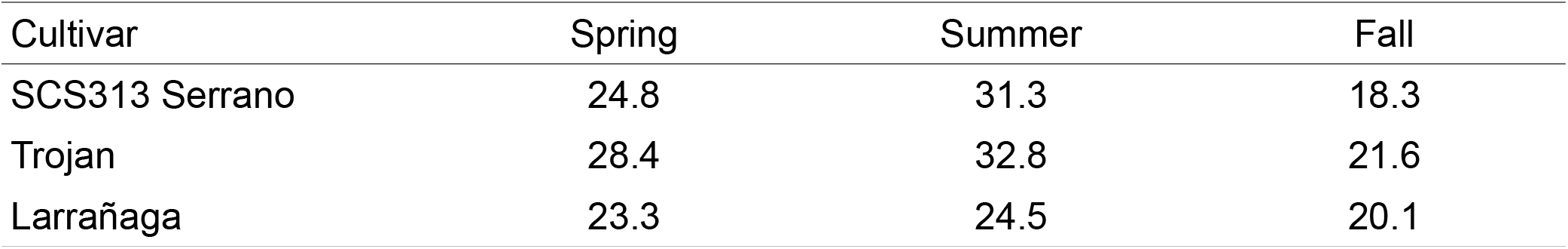
Season distribution for crude protein (CP) of the cultivars SCS313 Serrano, Trojan, and Larrañaga in Lages, SC, Brazil, from 2011 to 2013.

**Table 5.** Season distribution for in vitro dry matter digestibility (IVDMD) of the cultivars SCS313 Serrano, Trojan, and Larrañaga in Lages, SC, Brazil, from 2011 to 2013.

| Cultivar | Spring | Summer | Fall |
| --- | --- | --- | --- |
| SCS313 Serrano | 55 | 39.5 | 53.8 |
| Trojan | 61.6 | 46.5 | 50.1 |
| Larrañaga | 66.9 | 62.2 | 64.7 |

Like other lotus cultivars, Serrano has an adequate concentration of condensed tannins. The condensed tannins protect proteins from degradation and loss in the rumen, increasing intestinal flow and absorption at the same level, preventing bloating in cattle. The condensed tannins can also enable an acceptable performance in sheep even under high parasite levels (WAGHORN et al., 1994)

The challenges for commercial seed production of *L. uliginosus* include its indeterminate flowering with seed set over an extended period in summer and easy dehiscence of pods at maturity. Considering the shorter day length requirement of SCS313 Serrano, it has a high potential for commercial seed production; for production in low-latitude regions, such as southern Brazil, Uruguay, and eastern Australia; and for natural reseeding and enhance pasture persistence.

### Conclusions

The big trefoil cultivar SCS313 Serrano was selected for improving flowering ability and seed production in low-latitude regions at highlands of southern Brazil. This cultivar showed no reduction in dry matter yield when compared to the cultivars Trojan and Larrañaga, in the four locations evaluated. Thus, SCS313 Serrano is recommended to be used in mixtures with grass for pasture recovery in less intensive livestock operations, mainly on wet soils.

## Availability

The authors requested Plant Variety Protection for SCS313 Serrano in Brazil, Uruguay, Australia, and New Zealand. Seeds of SCS313 Serrano will be deposited at the USDA-ARS National Center for Genetic Resources Preservation, where it will be available for distribution after 20 years. This cultivar is exclusively licensed for commercial seed production and sales to PGG Wrightson Seeds.

